# Spike Reliability is Cell-Type Specific and Shapes Excitation and Inhibition in the Cortex

**DOI:** 10.1101/2024.06.05.597657

**Authors:** S. Russo, G. B. Stanley, F. Najafi

## Abstract

Neurons encode information in the highly variable spiking activity of neuronal populations, so that different repetitions of the same stimulus can generate action potentials that vary significantly in terms of the count and timing. How does spiking variability originate, and does it have a functional purpose? Leveraging the Allen Institute cell types dataset, we relate the spiking reliability of cortical neurons *in-vitro* during the intracellular injection of current resembling synaptic inputs to their morphologic, electrophysiologic, and transcriptomic classes. Our findings demonstrate that parvalbumin+ (PV) interneurons, a subclass of inhibitory neurons, show high reliability compared to other neuronal subclasses, particularly excitatory neurons. Through computational modeling, we predict that the high reliability of PV interneurons allows for strong and precise inhibition in downstream neurons, while the lower reliability of excitatory neurons allows for integrating multiple synaptic inputs leading to a spiking rate code. These findings illuminate how spiking variability in different neuronal classes affect information propagation in the brain, leading to precise inhibition and spiking rate codes.

## Introduction

Neurons encode continuously varying information through discrete action potentials. Action potentials have low reliability, so that neurons respond to multiple repetitions of the same stimulus with a different number of action potentials and at different timings^1– 7^. This leads to a large variability across repetitions, that is usually discarded through averaging while preserving the reliable neural response, invariant across repetitions.

Over the last decades, variability in neural activity has been progressively re-evaluated; from being a nuisance, it became a potentially valuable resource. Recent studies have postulated that variability in neuronal activity is essential for behaviors such as discrimination^8,9^, encodes important sensory and behavioral features^10–16^, and promotes efficient information coding^17,18^. The new understanding of the importance of neuronal reliability (and its opposite, variability) prompted further studies to identify its origins. These studies found that cortical responses to sensory stimuli are unreliable^19^ due to independent inputs, both from the environment or from the network^1,20–22^. One intriguing hypothesis is that multiple subtypes of cortical neurons may have distinct reliability properties, and their coordinated activity defines the overall reliability of the cortex in response to external stimuli^23^. Indeed, previous studies suggest that inhibitory neurons may respond more reliably to external stimuli compared to excitatory neurons^24,25^, thus suggesting that different cell types may exhibit different levels of reliability, but this has not been explicitly tested.

Unravelling reliability in distinct neuronal subtypes is crucial to understanding how they interact to process information in neural circuits. In this study, we set to address this question by investigating how spiking reliability encodes stimuli, through measures that incorporate both spike occurrence and temporal precision^26^. Specifically, we studied how reliability across action potentials induced by direct neuronal stimulation relates to cell-type specific neuronal properties, and how it impacts the effect of action potentials on target neurons.

## Results

First, we leveraged the Allen Institute cell types dataset^27^, which characterizes the morphologic, electrophysiologic, and transcriptomic features of mice and human cortical neurons in-vitro. Based on these features, we classified neurons in morphologic (aspiny, sparsely spiny, and spiny dendrites), electrophysiologic (fast and regular spiking; spike width below or above 400 μs, respectively), and transcriptomic cell-types (transgenic mouse lines).

To evaluate single-neuron spiking reliability, we analyzed patch-clamp recordings of neurons which had their synaptic inputs blocked (tissue bathed with 1 mM kynurenic acid and 0.1 mM picrotoxin) and which were directly stimulated with time-varying pink noise current, resembling physiological synaptic inputs^28,29^. To account for the different excitability across neurons, the intensity of the template stimulus was scaled as a function of the rheobase of each neuron, defined as the minimal electric current necessary to elicit an action potential for that neuron. This dataset contains two versions of noise (Noise 1 and 2) administered in current-clamp mode at different rheobase percentages: 75%, 100%, and 150%; (Figure 1A). Importantly, the time course and intensity of each noise was identical between sessions of the same neurons. This allowed us to assess reliability in the response to the same stimulus across repetitions. For this purpose, we analyzed only 1851 neurons (319 from humans, 1532 from mice) in which noise current stimulation was repeated more than once (2-8 times; Figure 1B: example mouse neuron; Figure 1C: all mouse neurons). To investigate reliability in this large sample of cells with a relatively low number of trials per cell, we designed specific measures invariant to the number of trials (Figure 2). To account for differences across species, mice and human neurons were analyzed separately (Figure 2 and Figure S1, respectively).

**Figure 1.**
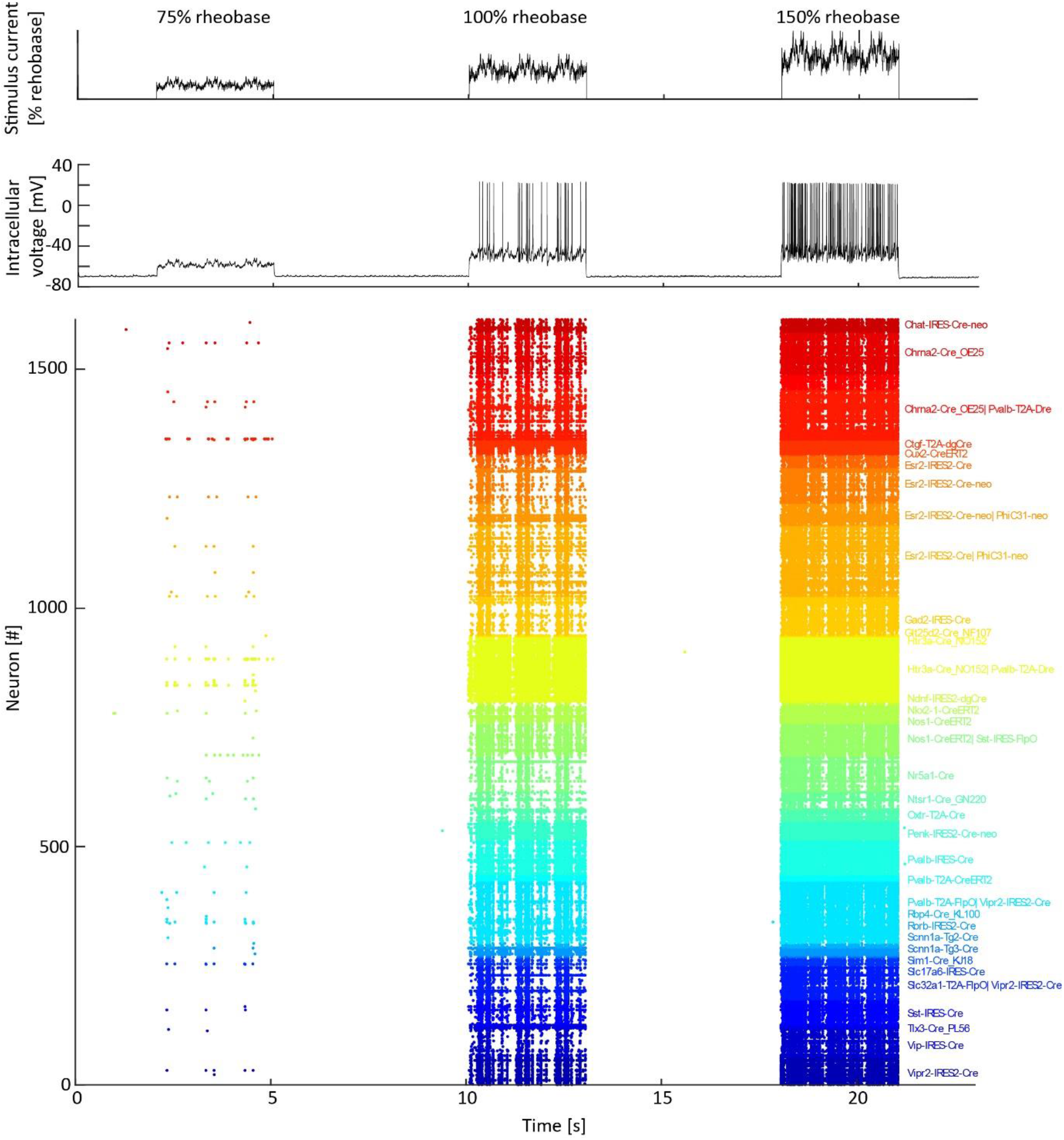
Pink noise current stimulation induces spiking activity in neocortical neurons in the Allen Institute cell types dataset. (Gouwens, Nat Neurosci, 2019). **A**. Time course of Noise 2 current (intensity: 75%, 100%, and 150% rheobase) administered in the stimulation protocol. **B**. Intracellular voltage of one recording from one representative neuron stimulated with Noise 2. **C**. Spike timing elicited by Noise 2 in each stimulated mouse neuron, color coded by transcriptomic cell type.

**Figure 2.**
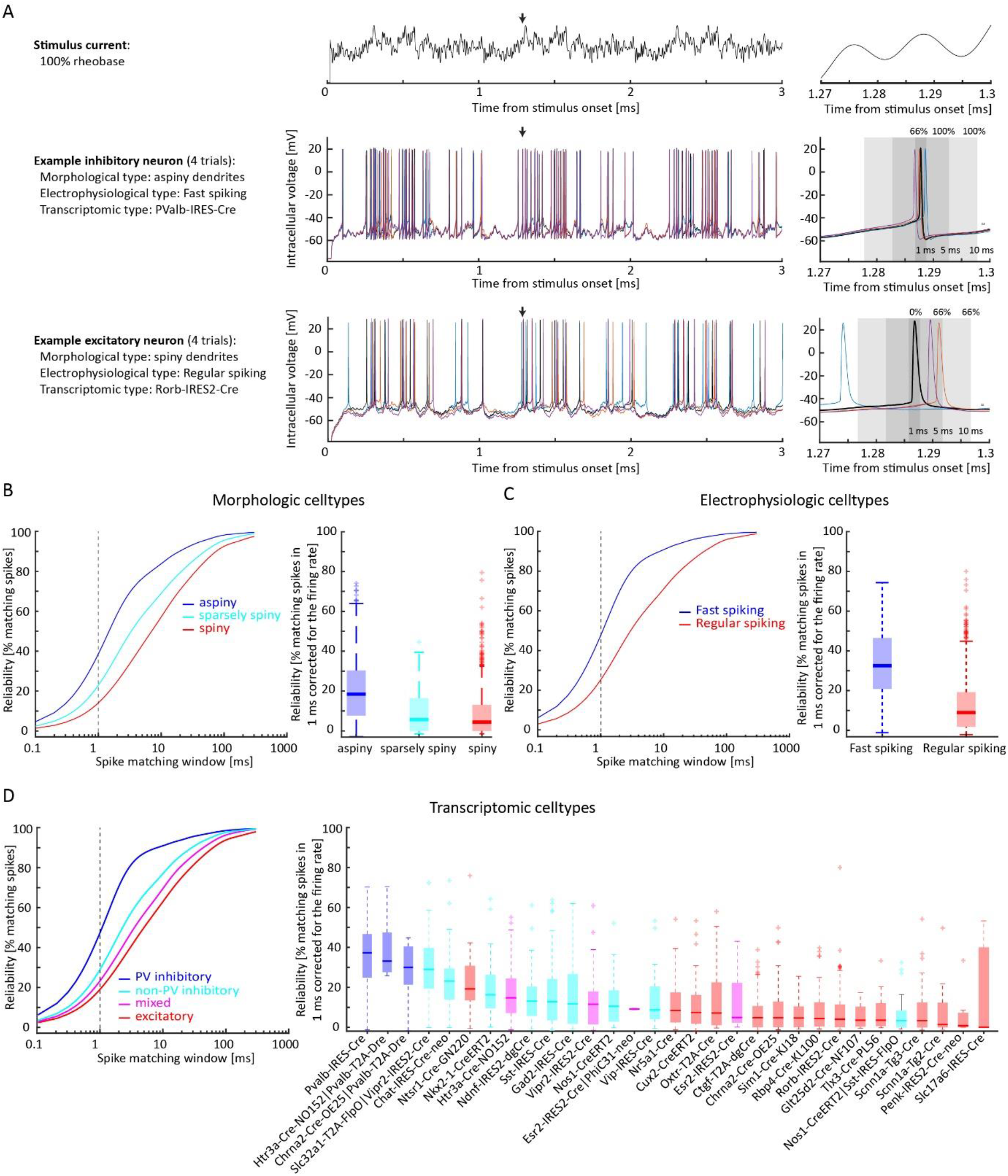
Noise stimulation induces more reliable response in morphologic, electrophysiologic, and transcriptomic cell-types associated with specific inhibitory neuronal populations in mice. **A**. Trace of the Noise 2 current at 100% rheobase (top). For one representative inhibitory and excitatory neuron (middle and bottom), we (left) summarize their morphologic, electrophysiologic, and transcriptomic cell-type, (center) show the intracellular voltage activity evoked by Noise 2 stimulation at 100% rheobase across 4 trials, and (right) show their reliability in the same time window (indicated by the black arrow in the central panel), as measured by the percentage of spikes falling within windows of increasing size from the selected spike. **B**. For morphologic cell types, we illustrate the probability density function of finding matching spikes across repetitions for increasing time windows (left; Noise 1 and 2 stimuli combined), together with the comparison of the reliability distribution across cell-types (right; Noise 1 and 2 stimuli combined ; window=1 ms, corrected for firing rate; boxplot reports median [thick line], 25° and 75° percentile [box], maximum and minimum within 1.5 interquartile range [whiskers], and outliers [+]). Color code indicates morphologic cell-types associated with excitatory (red) or inhibitory neurons (blue). **C**. Same representation as B for electrophysiologic cell-types. **D**. Same representation as B and C for transcriptomic cell-types grouped in PV (parvalbumin) inhibitory, non-PV inhibitory, mixed, and excitatory cell-types.

Figure 2A shows the activity evoked by four repetitions of noise stimulation in representative inhibitory and excitatory mouse neurons. We measured the percentage of matching spikes as the percentage of spikes in each repetition that occurred within a window of a given size with respect to the spikes in all other repetitions (Figure 2A, right). To quantify reliability, we assessed the percentage of matching spikes in a window of 1 ms corrected for firing rate (i.e. subtracting the same metric obtained after randomly shuffling spikes timings in 100 permutations).

In morphologic cell-types, we found that neurons with aspiny dendrites exhibit more reliable spiking activity compared to neurons with spiny and sparsely spiny dendrites (Figure 2B left). Indeed, we found that cells with aspiny dendrites show significantly higher reliability than neurons with spiny and sparsely spiny dendrites (Figure 2B right, 706 aspiny, 86 sparsely spiny, 740 spiny; Kruskal-wallis test; p<0.001; for details see Table S1).

Similarly, for electrophysiological cell-types, we found that fast spiking neurons show significantly higher reliability than regular spiking neurons (Figure 2C, 159 fast spiking, 1358 regular spiking; Kruskal-wallis test, p<0.001; for details see Table S1).

Importantly, aspiny dendrites and fast spiking activity are typical of PV inhibitory neurons, as opposed to the spiny dendrites and the regular spiking activity which are typical of excitatory neurons and non-PV inhibitory neurons^27^.

To further investigate this observation, we applied the same analysis to neurons categorized according to their transcriptomic cell-type, grouped in PV inhibitory neurons, non-PV inhibitory neurons, mixed neurons, and excitatory neurons^30^. We found that PV neurons show higher reliability than neurons in other transcriptomic cell-types (Figure 2D; 145 PV, 452 non-PV inhibitory, 172 mixed, and 763 excitatory neurons; Kruskal-Wallis test, p<0.001; for details see Table S1). For all categorizations, we verified that the stimulation at 150% rheobase intensity led to similar results.

To investigate the origins of spiking reliability, we measured its correlation with other neuronal properties. First, we quantified reliability of subthreshold fluctuations by measuring the correlation of the voltage recordings during subthreshold noise stimulations across repetitions (75% rheobase). While we found an overall high reliability in subthreshold potentials in all cell types^31^, we found a weak correlation between spiking reliability and subthreshold fluctuations’ reliability across neurons (Figure S2, r=0.11; p<0.001), suggesting that subthreshold reliability does not drive spiking reliability. Conversely, spiking reliability was highly correlated with other intrinsic neuronal features such as the membrane time constant (r=-0.61; p<0.001), rheobase (r=0.55; p<0.001), time to first spike evoked by a slow current ramp (r=0.49; p<0.001), and ratio between action potential peak upstroke and downstroke (r=-0.40; p<0.001). Altogether, these findings suggest that neuronal reliability may be shaped by the same neuronal properties that shape action potentials, such as voltage-dependent sodium and potassium channels, and cell surface area^32,33^.

Next, we studied spiking reliability in neurons derived from human samples. Similar to mouse neurons, we found that neurons with morphologic and electrophysiologic features associated with inhibitory PV neurons (i.e. aspiny and fast spiking, respectively) have higher spiking reliability; however, human neurons showed overall less reliable spiking activity than mice neurons (Figure S1, 2-way ANOVA p<0.001).

To understand the network consequence of our findings, we modelled the impact of reliability on target neurons. To this aim, we built a model specifically designed for testing the effect of input reliability^26,34^, in which one Izhikevich spiking neuron^35^ receives gaussian synaptic inputs from a population of either excitatory or inhibitory neurons, that we parametrized in terms of timing variability (Figure 3A, B) and number of neurons (Figure 3C). Inhibitory inputs with low variability resulted in strong and sharp hyperpolarization, while highly variable inhibitory inputs led to weak prolonged hyperpolarization (Figure 3A). Indeed, the strength of the hyperpolarization, computed as the difference between the baseline voltage and the minimum voltage, progressively declined with increasing levels of variability (Figure 3A).

**Figure 3.**
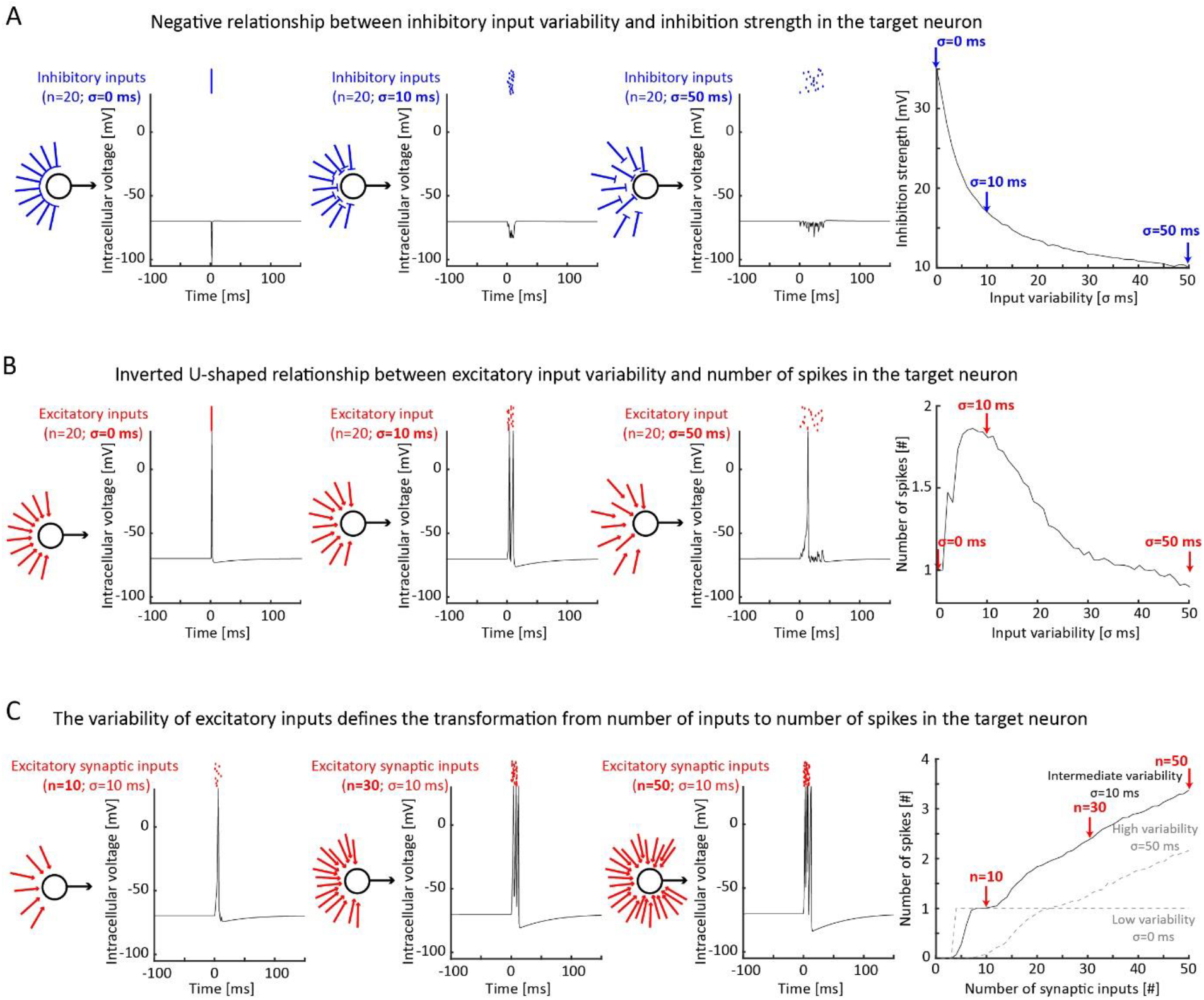
Computational modeling shows that high reliability of inhibitory inputs increases the strength of the inhibition in target neurons while intermediate variability of excitatory inputs increases the number of spikes generated in target neurons. **A**. From left to right: Schematic of the model showing the administration of inhibitory inputs to the target neuron and intracellular voltage evoked in the target neuron by 20 inhibitory inputs with low (left), intermediate (center), and high (right) levels of variability (shown by variability in input position). Plot showing the depth of the inhibition strength as a function of input reliability (500 iterations; representative cases pointed by blue arrows). Highly reliable inhibitory inputs generate a sharp, intense hyperpolarization in the target neuron, while less reliable inhibitory inputs generate more prolonged and superficial hyperpolarization. **B**. Same as A for 20 excitatory inputs with low (left; σ = 0 ms), intermediate (center; σ = 10 ms), and high (right; σ = 50 ms) levels of variability. Plot showing the number of spikes in the target neuron as a function of input reliability (500 iterations; representative cases pointed by red arrows). Excitatory inputs with intermediate variability generate more spikes than those with low and high variability. **C**. From left to right: schematic and intracellular potential evoked by the administration of 10 (left), 40 (center), and 50 (right) excitatory inputs with intermediate variability (σ = 10 ms), respectively. Plot showing the number of spikes as a function of the number of excitatory synaptic inputs for intermediate (solid black line; representative cases pointed by red arrows; 500 iterations), low, and high (dashed grey lines) levels of reliability.

Excitatory inputs with very low or very high variability led to one single spike in the target neuron. In contrast, intermediate levels of excitatory input variability led to the largest number of spikes in the target neuron (Figure 3B). This is captured by an inverse u shape relationship between excitatory input variability and number of spikes. Next, we varied the number of excitatory inputs in our model (Figure 3C). We found that excitatory inputs without any variability always generated one single spike in the target neuron, once that the number of excitatory inputs was sufficient to generate one spike. In contrast, temporally variable excitatory inputs led to increasingly more spikes in the target neuron, proportional to the number of excitatory inputs. Importantly, the variability of the inputs determined the steepness of the relationship between number of inputs and number of spikes. Altogether, our model suggest that low variability maximizes the impact of inhibitory inputs on their target neuron; conversely, moderate degrees of variability, defined by synaptic and neuronal properties, can maximize the impact of excitatory inputs on their target neuron.

## Discussion

Our findings indicate that action potential reliability in the timescale of few milliseconds is a cell-type specific feature related to the intrinsic properties of each neuron. Specifically, we found that PV neurons exhibit high reliability with respect to other cell types, particularly to excitatory neurons (Figure 2). Furthermore, our model predicted that the low variability of inhibitory neurons allows for strong inhibition, while the high variability of excitatory neurons allows for rate coding in downstream neurons (Figure 3).

To our knowledge, our study is the first that examines neuronal spiking reliability of different cell types following direct stimulation, while previous studies examined neuronal spiking reliability following indirect stimulation, either by stimulating pre-synaptic excitatory neurons^25,36^ or by presenting sensory inputs to animals^24^. While our findings are aligned with these previous studies which suggest high temporal precision in the spiking activity for inhibitory neurons, our study suggests that this feature of inhibitory neurons emerges also from intrinsic properties, and not necessarily from synaptic mechanisms. It is noteworthy that while our study was performed *in vitro*, our finding that PV neurons exhibit higher reliability than other cell types aligns with previous *in vivo* findings ^24,25^.

We have demonstrated that among inhibitory neurons, high reliability is typical of PV, but not SST (somatostatin) and VIP (vasointestinal peptide) neurons. This is in line with previous studies showing the distinct role of PV neurons in increasing reliability in responses to visual stimuli^37^ and in simulated circuits^38^. The highly synchronous spiking in PV neurons can explain their role in enhancing signal, reducing noise^23,37,39^, as well as functionally sculpting neuronal ensembles in the cortex^40^. It also highlights their role in cortical discrimination of auditory^41^, visual^37^, and somatosensory stimuli^42^. The reliability properties of PV, SST, and VIP neurons suggest that they may play different functional roles. For example, among inhibitory neurons, PV neurons may gate incoming inputs by providing a timely, synchronous inhibition, while SST and VIP neurons may downmodulate cortical excitability through prolonged inhibition.

Contrary to PV inhibitory neurons, excitatory neurons exhibit variability in their spiking activity. The reliability differences between excitatory, PV, SST, and VIP neurons suggest that different cell types may encode stimulus information through different algorithms^43^, with different functional significances. The variability of excitatory neurons is functionally significant as it facilitates the generation of rate codes within the cortex. Our model indicates that without this temporal variability in spiking activity, excitatory neurons would elicit a consistent firing rate in response to synchronous inputs, regardless of the number of excitatory inputs received (see Figure 3C).

Our work demonstrates that spiking reliability emerges, at least in part, from intrinsic cellular properties, and that it varies among distinct neuronal cell types underlying specific functions in excitatory and inhibitory neurons. Our findings provide insight into the circuit and behavioral consequences of reliability in distinct neuronal cell types. Further studies can shed light on how reliability emerges and what is its functional significance. In particular, studies at molecular scale will illuminate how membrane properties and specific ionic channels shape the reliability of different neurons and neuronal compartments; *in-vitro* studies without synaptic blockers and *in-vivo* studies will illuminate the network consequences of different reliability levels in distinct cell types.

## Supporting information

Statistical Table S1

## Bibliography

1. Nolte, M., Reimann, M. W., King, J. G., Markram, H. & Muller, E. B. Cortical reliability amid noise and chaos. Nat. Commun. 10, 3792 (2019).

2. Victor, J. D. & Purpura, K. P. Nature and precision of temporal coding in visual cortex: a metric-space analysis. J. Neurophysiol. 76, 1310–1326 (1996).

3. Tolhurst, D. J., Movshon, J. A. & Dean, A. F. The statistical reliability of signals in single neurons in cat and monkey visual cortex. Vision Res. 23, 775–785 (1983).

4. Butts, D. A. et al. Temporal precision in the neural code and the timescales of natural vision. Nature 449, 92–95 (2007).

5. Bair, W. & Koch, C. Temporal Precision of Spike Trains in Extrastriate Cortex of the Behaving Macaque Monkey. Neural Comput. 8, 1185–1202 (1996).

6. Liu, R. C., Tzonev, S., Rebrik, S. & Miller, K. D. Variability and Information in a Neural Code of the Cat Lateral Geniculate Nucleus. J. Neurophysiol. 86, 2789–2806 (2001).

7. Reich, D. S., Victor, J. D., Knight, B. W., Ozaki, T. & Kaplan, E. Response Variability and Timing Precision of Neuronal Spike Trains In Vivo. J. Neurophysiol. 77, 2836–2841 (1997).

8. Waschke, L., Kloosterman, N. A., Obleser, J. & Garrett, D. D. Behavior needs neural variability. Neuron 109, 751–766 (2021).

9. Arazi, A., Censor, N. & Dinstein, I. Neural Variability Quenching Predicts Individual Perceptual Abilities. J. Neurosci. 37, 97–109 (2017).

10. Hussar, C. & Pasternak, T. Trial-to-trial variability of the prefrontal neurons reveals the nature of their engagement in a motion discrimination task. Proc. Natl. Acad. Sci. 107, 21842–21847 (2010).

11. Festa, D., Aschner, A., Davila, A., Kohn, A. & Coen-Cagli, R. Neuronal variability reflects probabilistic inference tuned to natural image statistics. Nat. Commun. 12, 3635 (2021).

12. Reinagel, P. & Reid, R. C. Temporal coding of visual information in the thalamus. J. Neurosci. Off. J. Soc. Neurosci. 20, 5392–5400 (2000).

13. Reich, D. S., Mechler, F., Purpura, K. P. & Victor, J. D. Interspike Intervals, Receptive Fields, and Information Encoding in Primary Visual Cortex. J. Neurosci. 20, 1964–1974 (2000).

14. Panzeri, S., Brunel, N., Logothetis, N. K. & Kayser, C. Sensory neural codes using multiplexed temporal scales. Trends Neurosci. 33, 111–120 (2010).

15. Nicola, W., Newton, T. R. & Clopath, C. The impact of spike timing precision and spike emission reliability on decoding accuracy. Sci. Rep. 14, 10536 (2024).

16. Kasamatsu, T., Polat, U., Pettet, M. W. & Norcia, A. M. Colinear facilitation promotes reliability of single-cell responses in cat striate cortex. Exp. Brain Res. 138, 163–172 (2001).

17. Scaglione, A., Moxon, K. A., Aguilar, J. & Foffani, G. Trial-to-trial variability in the responses of neurons carries information about stimulus location in the rat whisker thalamus. Proc. Natl. Acad. Sci. 108, 14956–14961 (2011).

18. Stein, R. B., Gossen, E. R. & Jones, K. E. Neuronal variability: noise or part of the signal? Nat. Rev. Neurosci. 6, 389–397 (2005).

19. Kara, P., Reinagel, P. & Reid, R. C. Low response variability in simultaneously recorded retinal, thalamic, and cortical neurons. Neuron 27, 635–646 (2000).

20. Faisal, A. A., Selen, L. P. J. & Wolpert, D. M. Noise in the nervous system. Nat. Rev. Neurosci. 9, 292–303 (2008).

21. Rusakov, D. A., Savtchenko, L. P. & Latham, P. E. Noisy Synaptic Conductance: Bug or a Feature? Trends Neurosci. 43, 363–372 (2020).

22. Gur, M., Beylin, A. & Snodderly, D. M. Response variability of neurons in primary visual cortex (V1) of alert monkeys. J. Neurosci. Off. J. Soc. Neurosci. 17, 2914–2920 (1997).

23. Wehr, M. & Zador, A. M. Balanced inhibition underlies tuning and sharpens spike timing in auditory cortex. Nature 426, 442–446 (2003).

24. Tao, C. et al. Synaptic Basis for the Generation of Response Variation in Auditory Cortex. Sci. Rep. 6, 31024 (2016).

25. Levi, A., Spivak, L., Sloin, H. E., Someck, S. & Stark, E. Error correction and improved precision of spike timing in converging cortical networks. Cell Rep. 40, 111383 (2022).

26. Wang, H.-P., Spencer, D., Fellous, J.-M. & Sejnowski, T. J. Synchrony of thalamocortical inputs maximizes cortical reliability. Science 328, 106–109 (2010).

27. Gouwens, N. W. et al. Classification of electrophysiological and morphological neuron types in the mouse visual cortex. Nat. Neurosci. 22, 1182–1195 (2019).

28. de Ruyter van Steveninck, R. R., Lewen, G. D., Strong, S. P., Koberle, R. & Bialek, W. Reproducibility and Variability in Neural Spike Trains. Science 275, 1805–1808 (1997).

29. Mainen, Z. F. & Sejnowski, T. J. Reliability of spike timing in neocortical neurons. Science 268, 1503–1506 (1995).

30. Tasic, B. et al. Shared and distinct transcriptomic cell types across neocortical areas. Nature 563, 72–78 (2018).

31. Carandini, M. Amplification of Trial-to-Trial Response Variability by Neurons in Visual Cortex. PLOS Biol. 2, e264 (2004).

32. Golowasch, J. et al. Membrane capacitance measurements revisited: dependence of capacitance value on measurement method in nonisopotential neurons. J. Neurophysiol. 102, 2161–2175 (2009).

33. Bean, B. P. The action potential in mammalian central neurons. Nat. Rev. Neurosci. 8, 451–465 (2007).

34. Wright, N. C. et al. Rapid Cortical Adaptation and the Role of Thalamic Synchrony during Wakefulness. J. Neurosci. Off. J. Soc. Neurosci. 41, 5421–5439 (2021).

35. Izhikevich, E. M. Simple model of spiking neurons. IEEE Trans. Neural Netw. 14, 1569–1572 (2003).

36. Jouhanneau, J.-S., Kremkow, J. & Poulet, J. F. A. Single synaptic inputs drive high-precision action potentials in parvalbumin expressing GABA-ergic cortical neurons in vivo. Nat. Commun. 9, 1540 (2018).

37. Zhu, Y., Qiao, W., Liu, K., Zhong, H. & Yao, H. Control of response reliability by parvalbumin-expressing interneurons in visual cortex. Nat. Commun. 6, 6802 (2015).

38. Guo, L. & Kumar, A. Role of interneuron subtypes in controlling trial-by-trial output variability in the neocortex. Commun. Biol. 6, 1–13 (2023).

39. Cardin, J. A. et al. Driving fast-spiking cells induces gamma rhythm and controls sensory responses. Nature 459, 663–667 (2009).

40. Agetsuma, M., Hamm, J. P., Tao, K., Fujisawa, S. & Yuste, R. Parvalbumin-Positive Interneurons Regulate Neuronal Ensembles in Visual Cortex. Cereb. Cortex N. Y. N 1991 28, 1831–1845 (2018).

41. Nocon, J. C. et al. Parvalbumin neurons enhance temporal coding and reduce cortical noise in complex auditory scenes. Commun. Biol. 6, 751 (2023).

42. Yeganeh, F. et al. Effects of optogenetic inhibition of a small fraction of parvalbumin-positive interneurons on the representation of sensory stimuli in mouse barrel cortex. Sci. Rep. 12, 19419 (2022).

43. Kumbhani, R. D., Nolt, M. J. & Palmer, L. A. Precision, reliability, and information-theoretic analysis of visual thalamocortical neurons. J. Neurophysiol. 98, 2647–2663 (2007).

