## Supplemental materials for "Spike Reliability is Cell-Type Specific and Shapes Excitation and Inhibition in the Cortex"

### Supplementary figures

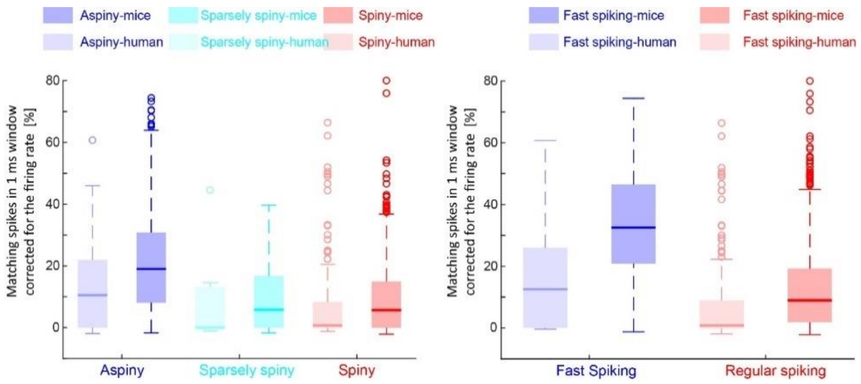


**Figure S1. Reliability across cell types and species.** Matching spikes in 1 ms window corrected for the firing rate for morphologic (left) and electrophysiologic (right) cell types in human (shaded boxes) and mice samples (solid boxes). Boxplot reports median [thick line], 25° and 75° percentile [box], maximum and minimum within 1.5 interquartile range [whiskers], and outliers [o].

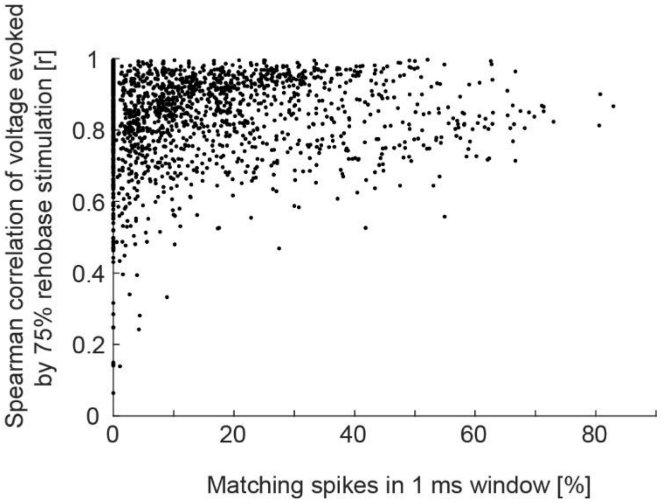


**Figure S2. Relationship between spike reliability and subthreshold reliability.** Scatter plot showing the relationship between the reliability of subthreshold fluctuations, computed as the average spearman correlation across repetitions of the noise stimulation at 75% rheobase intensity, and the percentage of matching spikes in 1 ms window.

### Supplementary table

**Table S1**

The statistical table file reports the results of the Kruskal-Wallis tests and 2-way ANOVA tests. The title of the section referring to each test is highlighted in grey. For easiness of read, the names of the transcriptomic cell types are reported in the orange column beside the relative Kruskal-Wallis test.

### Supplementary methods

Metadata

Data was obtained from the Allen institute cell types datasets. Spike timings, transcriptomic cell types, and morphologic cell types, were extracted from the original dataset. Spike width was computed as follows. First, we averaged all the spikes evoked during the noise stimulation in one neuron. Using the so obtained average spike, we computed the spike width for that neuron as the duration of the spike at half of the spike amplitude.

### Spike reliability analysis

To compute the cumulative probability density, we computed the latency between each spike of one neuron and the closest spike across all the sessions of the same neuron. These values were aggregated across sessions and neurons. Using the obtained distribution, we computed the relative cumulative density function using the ksdensity from Matlab (function parameter: cdf).

To compute the matching spikes in 1 ms window, we computed for each neuron the percentage of spikes occurring within 1 ms from at least one spike in each other trial. To control for the firing rate, we subtracted to this value the same metric obtained from shuffled data, in which the number of spikes in each session was kept constant, but their latency was randomly permuted within the stimulus time window.

We compared different cell types within mice using a Kruskal-Wallis test. Comparisons across cell types and species we carried out using a two-way ANOVA test.

### Computational model

The model was constituted by a set of synaptic inputs – parametrized in terms of number, variability, and polarity – projecting into a target neuron. Each synaptic input was modelled as a gaussian current, with fixed amplitude and duration across neurons. The polarity of the current was determined by the input type, being negative (hyperpolarizing) for inhibitory inputs and positive (depolarizing) for excitatory inputs. To introduce variability in inputs' latencies, the timing of each input was randomly sampled from a gaussian distribution with standard deviation = $\sigma$.

The target neuron was modelled as a single-compartment Izhikevich neuron in spiking regimen (voltage capped to 30 mV).
